# Decoding *RAP1*’s Role in Yeast mRNA Splicing

**DOI:** 10.1101/2025.04.04.647307

**Authors:** Siddhant Kalra, Joseph D. Coolon

## Abstract

Messenger RNA (mRNA) splicing is a fundamental and tightly regulated process in eukaryotes, where the spliceosome removes non-coding sequences from pre-mRNA to produce mature mRNA for protein translation. Alternative splicing enables the generation of multiple RNA isoforms and protein products from a single gene, regulating both isoform diversity and abundance. While splicing is widespread in eukaryotes, only ∼3% of genes in *Saccharomyces cerevisiae* undergo splicing, with most containing a single intron. However, intron-containing genes, primarily ribosomal protein genes, are highly expressed and constitute about one-third of the total mRNA pool. These genes are transcriptionally regulated by Repressor Activator Protein 1 (*RAP1*), prompting us to investigate whether *RAP1* influences mRNA splicing. Using RNA sequencing, we identified a novel role for *RAP1* in alternative splicing, particularly in intron retention (IR) while minor effects were observed on alternative 3’ and 5’ splice site usage. Many IR-containing transcripts introduced premature termination codons, likely leading to degradation via nonsense-mediated decay (NMD). Consistent with previous literature, genes with predicted NMD in our study also had reduced overall expression levels suggesting that *RAP1* plays an important role in this understudied mechanism of gene expression regulation.

## Introduction

Eukaryotic gene expression of protein coding genes is a complex and highly regulated multi-step process. First, specific DNA sequences throughout the genome, typically specified by the combinatorial binding of regulatory proteins, are transcribed to produce RNAs. These RNAs undergo co-transcriptional RNA processing to produce mature messenger RNA (mRNA). Once formed mRNA can then be exported from the nucleus to the cytoplasm where it can be translated into protein by the ribosome (1). Critical to this process is the removal of non-coding sequences present in nascent RNA transcripts by the spliceosome protein complex, splicing together exons to generate an RNA with a long open reading frame (ORF) of coding sequence optimized for subsequent translation (2–4).

Almost all eukaryotes have introns and share common mechanisms of mRNA splicing indicating an ancient origin and its persistence across deep evolutionary time demonstrates its essential role in the regulation of gene expression (5,6). Because mRNA splicing can lead to the creation of different combinations of sequences from a single gene, so-called alternative splicing (AS) produces alternative mRNA isoforms that encode for different proteins with different functions. This serves to massively increase the coding potential for the genome as well as proteome complexity and diversity (5) which has allowed for further fine-tuning the functionality and specificity of the respective proteins made for their distinct roles in the organism (7–9). Additionally, the efficiency of this process directly influences the overall levels of gene expression (10–12) where introns have been demonstrated to enhance both transcription and translational output for genes where they occur (13). Moreover, in humans, there is a positive correlation between mRNA stability and the presence of introns within a gene leading to a phenomenon where genes containing introns exhibit greater mRNA stability compared to their intronless counterparts (14,15).

Alternative splicing can take many forms, including Exon Skipping (ES, sometimes called Cassette Exon), Mutually Exclusive exons (ME), Intron Retention (IR), Alternative 3’ Splice Site (A3SS) and Alternative 5’ Splice Site (A5SS) splicing events. Both ES and ME require at least 2 introns while IR, A3SS and A5SS only require 1 intron per gene (2,16). Moreover, at each intron, different types of splicing events can occur further increasing the number of potential isoforms produced by a gene. Mutations occurring within exons, introns, or other influencing factors can disrupt the splicing machinery, potentially leading to the generation of aberrant transcripts (17– 20) with alternative open reading frames that can have profound impacts on phenotypes including generation of human disease states (21–23). For example, a mutation within the branch point sequence of the human Neurofibromatosis type 1 (*NF1)* gene can give rise to a partial retention of intron 15, leading to Neurofibromatosis type 1 (24) and a point mutation in the splice donor site of the Major Intrinsic Protein Of Lens Fiber (*MIP)* gene forcing an exon skipping event that gives rise to cataracts (25).

In plants, fungi, and unicellular eukaryotes, IR is the most frequent type of alternative splicing observed (26–28) and multiple studies have provided evidence that IR in humans is essential for gene expression regulation during development (29,30). Similarly, in *Drosophila* IR facilitates X chromosome dosage compensation and is essential for proper gene regulation (31,32). When introns are retained in mature mRNA transcripts, they can encode new protein variants with the additional amino acid sequences derived from the intron, resulting in altered or novel functions compared to proteins produced when introns are spliced out. Sometimes this can lead to protein-coding isoforms associated with disease conditions including Alzheimer’s disease (33), prostate cancer (34) and lymphoma (35).

Transcripts with IR can sometimes be retained within the nucleus, where they either await splicing signals for further processing or are targeted for degradation (27). In certain cases, IR can lead to the inclusion of a premature termination codon (PTC) (27) that will produce truncated proteins potentially detrimental to cellular functions that contribute to the development of diseases. For example, the retention of a 192 bp segment of intron 13 in the gene encoding pyruvate carboxylase introduces a PTC, resulting in pyruvate carboxylase deficiency and subsequent delayed neurodevelopment (36). Similarly, intron retention in human *EIF2B5* gene leads to the production of a truncated protein, which has been implicated as a contributing factor in head and neck cancers (29,37).

In other cases, intron-retained transcripts containing PTC are directed to degradation via the nonsense-mediated decay (NMD) pathway (27,28,38,39). This pathway functions as a quality control mechanism for post-transcriptional regulation and is found in all eukaryotes. IR coupled with NMD is often associated with a reduction in gene and protein expression by degrading mRNA transcripts containing PTCs. Thus, NMD helps the cell to control both the abundance and quality of mRNAs, contributing to overall gene regulation and protein abundance (40,41) impacting both normal physiology and disease states. For example, IR followed by NMD fine-tunes gene expression levels during human granulocyte differentiation (42) and can also locally regulate mRNA stability and help neuronal axon guidance (43). Similarly, fungi can modulate the levels of cellulase transcripts in response to altered environmental conditions using IR followed by NMD ultimately controlling enzyme abundance in (28). In contrast to the regulatory roles described above, IR followed by NMD is also associated with both lung cancer (44) and hepatocellular carcinoma (45).

In *Saccharomyces cerevisiae* a relatively small proportion of genes contain introns (∼340 genes out of ∼7000 genes), with almost all having a single intron (11,12,46,47), however, due to the high expression levels of these genes, the majority of mRNA transcripts generated by a cell (∼70%) originate from them (11,46,48). A substantial proportion (∼90%) of these transcripts are dedicated to the expression of ribosomal protein genes (RPGs) (46,48,49). The presence of introns in RPGs contributes to the survival of yeast under stressful conditions, thereby conferring a fitness advantage for organismal growth when present (47,50–53). Eliminating introns from RPGs influences yeast survival and growth demonstrating their important role in RPG function and/or regulation. It has been observed that introns have the capacity to modulate the expression of RPGs, either enhancing or attenuating gene expression through intra or intergenic intron-dependent mechanisms (46,54,55). Furthermore, yeast retained introns containing PTCs also result in targeting for NMD causing reduced overall expression demonstrating a lesser-known mechanism for gene repression (56–59)

In yeast over 70% of ribosomal protein genes contain at least one intron (12,46). Notably, the essential yeast transcription factor Repressor Activator Protein 1 (*RAP1*) regulates approximately 5% of the yeast genome (60–62). Its primary targets include telomere-proximal genes, homothallic mating (HM) loci, glycolytic genes, DNA repair genes, and most importantly for this study, ribosomal protein genes (60,63–70). Our previous study utilized the Tet-off system to modulate the expression levels of *RAP1* in the presence of the doxycycline (Dox) in Yeast Dextrose Peptone (YPD) growth media (62). This design ensured an inverse relationship between the concentration of Dox and the expression level of *RAP1*. Our findings indicated a significant influence of *RAP1* reduction on the expression levels of RPGs. We found that RPGs expression was significantly reduced in response to reduction in *RAP1* levels suggesting that *RAP1* plays an activating role in their regulation (62,71). Given that ribosomal protein genes are intron-containing genes and are regulated by *RAP1*, our investigation aims to determine whether alterations in *RAP1* levels impact alternative splicing, a function that to our knowledge has not been previously demonstrated for *RAP1*.

The human ortholog of yeast *RAP1*, called Telomeric Repeat-Binding Factor 2-Interacting Protein 1 (*TERF2IP*), is a multifunctional protein that plays a critical role in numerous processes, strikingly these are the same processes regulated by yeast *RAP1. TERF2IP* is critical for maintaining telomeric DNA integrity and chromosome stability as part of the shelterin complex (72–75). In addition to its telomeric functions, *TERF2IP* acts as a transcription factor involved in regulating metabolic pathways and non-canonical NF-κB signaling (72,76–78). Its dysregulation has been implicated in various cancers, including breast (72,79), gastric (72,80), colorectal (72,81), renal (72,82), and non-small cell lung carcinoma (72,83), where it is often overexpressed. Conversely, reduced *TERF2IP* expression has been associated with papillary thyroid cancer (84). *TERF2IP* is also considered a predictive marker for evaluating chemotherapy success in breast (85) and colon cancer (81). Understanding *RAP1’s* role in yeast RNA splicing should provide valuable insights into whether *TERF2IP* also has roles in splicing which could further elucidate its broader implications in processes like cancer development and gene regulation.

## Results

### Testing for roles of *RAP1* in regulating alternative splicing

To determine if the well-studied essential yeast transcriptional regulator *RAP1* has previously uncharacterized functions in mRNA splicing, we used the Tet-off system (62,86) to sequentially titrate the expression level of *RAP1* by varying levels of Dox in growth media and estimated the effects that altered *RAP1* abundance had on alternative splicing genome-wide using RNA-seq (Fig. 1A). Data generated was subjected to a quality check with FASTQC (87), alignment to the yeast genome with the STAR aligner (88) and alternative splicing was quantified with the SpliceWiz (89) package in R (Fig. 1B).

**Fig 1:**
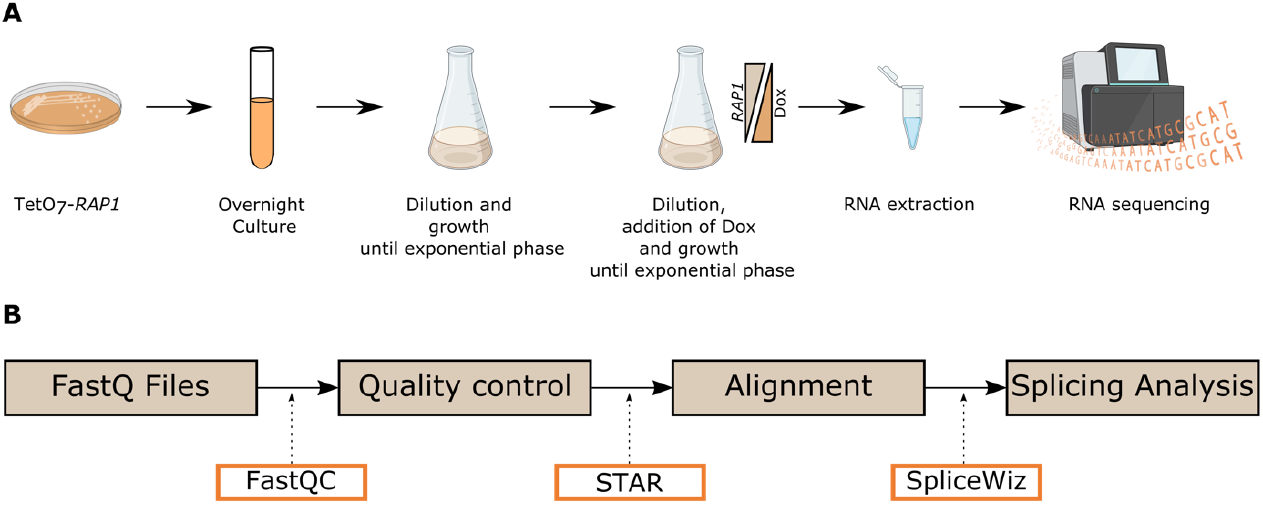
Workflow. A) Experimental workflow illustrating the sample preparation process for RNA extractions. B) Bioinformatic pipeline for downstream analysis of RNA-seq data. Tools and algorithms employed at each stage are also highlighted.

We first analyzed our control samples (zero Dox in media, maximum *RAP1* levels) with the splicing pipeline outlined in Fig. 1B and observed a total of 219 genes (out of 340 total intron containing genes) contributing to 229 Intron Retention (IR) splicing events (one gene can have different introns or locations where IR occurs hence contributing to different splicing events) (Fig. 2A), 23 genes contributing to 25 Alternative 3’ Splice Site (A3SS) splicing events (Fig. 2A), and 7 genes contributing to 7 Alternative 5’ Splice Site (A5SS) splicing events (Fig. 2A)(S-Table 1). The observed abundance of IR in control cells grown in standard conditions is consistent with the estimates found in prior literature (47,53,90).

**Fig 2:**
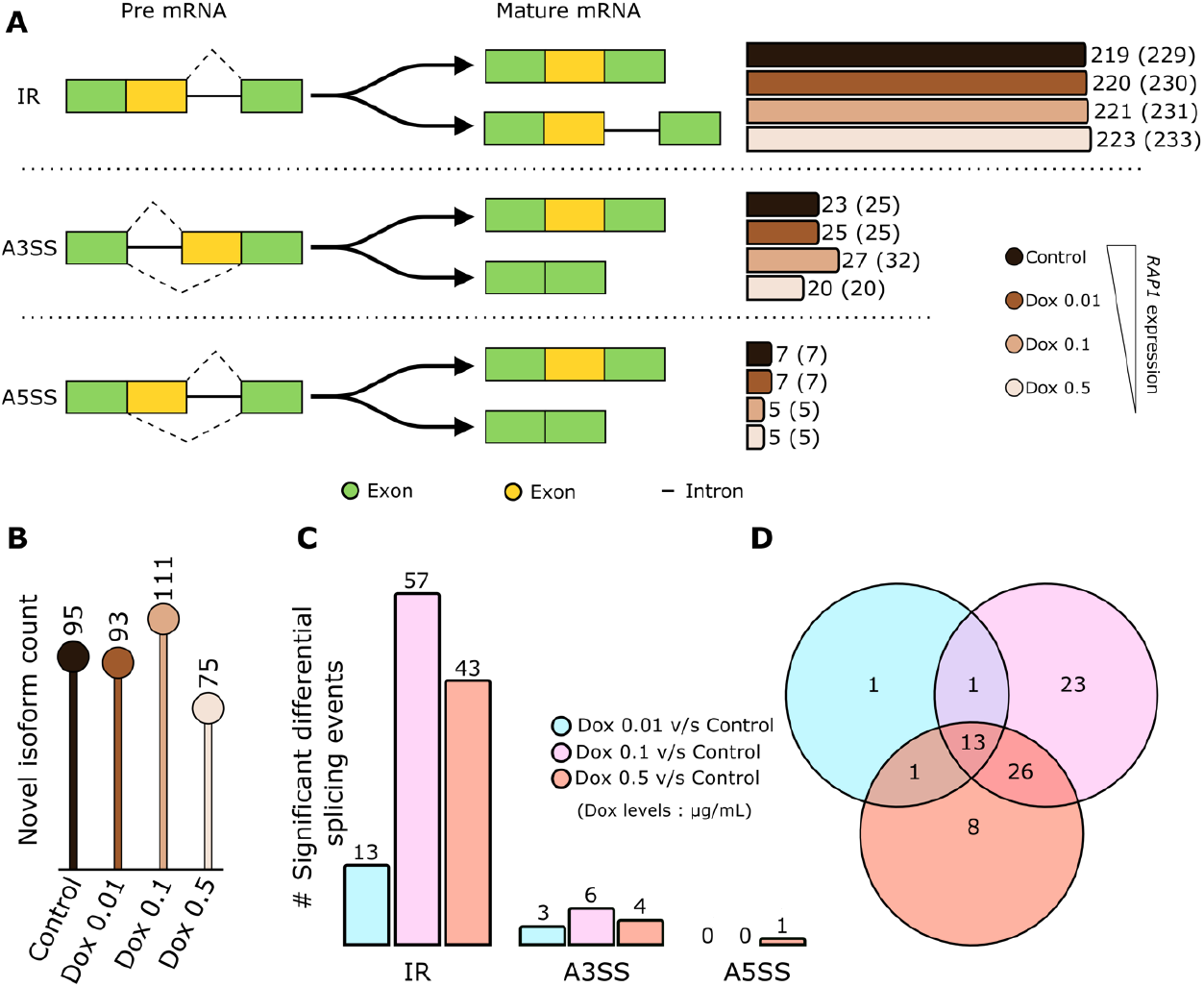
Quantification of splicing events. A) Quantification of different splicing events observed. The green and yellow color on the left indicates exons while the solid black line indicates the presence of an intron. The dashed black line represents alternative splicing junctions. On the right, different shades of brown represent different *RAP1* levels followed by the quantification of genes while the quantification of splicing events is mentioned in parentheses. B) Determination of novel isoforms produced at different (*RAP1*) Dox levels. C) Bar plots indicating the quantification of significant differential splicing events at various *RAP1* levels (compared to control). D) Venn diagram depicting the overlap of significant splicing events across different comparisons.

Next, we wanted to identify possible AS events at various reduced levels of *RAP1* expression. We found an increasing trend wherein the number of genes with IR increased as *RAP1* levels decreased (Fig. 2A). At a Dox concentration of 0.01 µg/mL we observed 230 IR events (from 220 genes), 231 IR events were observed (from 221 genes) at Dox 0.1 µg/mL and 233 IR events observed (from 223 genes) at Dox 0.5 µg/mL (Fig. 2A) (S-Table 1). We did not find a relationship for A3SS as *RAP1* levels were reduced. We observed 25 A3SS events (from 25 genes) at Dox 0.01 µg/mL, 32 A3SS events (from 27 genes) at Dox 0.1 µg/mL and 20 A3SS events (from 20 genes) at Dox 0.5 µg/mL (Fig. 2A) (S-Table 1). We again found no consistent trend for A5SS splicing events as *RAP1* expression was reduced. We identified 7 A5SS events at Dox 0.01 µg/mL while 5 events were observed at Dox 0.1 µg/mL and Dox 0.5 µg/mL (Fig. 2A) (S-Table 1).

We also determined whether and how many novel isoforms were produced in response to altered *RAP1* abundance. At Dox 0, we observed the generation of 95 novel transcripts (not currently annotated in genome-wide yeast databases), accounting for 2.67% of the total transcripts produced under this condition (Fig. 2B). At Dox 0.01, 93 novel transcripts were identified, contributing 2.57% of the total transcripts generated at this level (Fig. 2B). At Dox 0.1, the number of novel transcripts increased to 111, representing 3.02% of the total transcripts. In contrast, at Dox 0.5, 75 novel transcripts were generated, contributing 2.11% to the total transcripts (Fig. 2B).

### Quantification of significantly differentially spliced events following *RAP1* titration

We next asked how many of the observed events were significantly differently spliced in comparison to significantly differentially spliced our control condition (0 Dox). When comparing Dox 0.01 to the control, our analysis identified 13 significantly differentially spliced IR events and 3 A3SS significantly differentially spliced events (Fig. 2C) (S-Table 2). At Dox 0.1 versus control, we observed 57 significantly different IR events and 6 significantly different A3SS events (Fig. 2C) (S-Table 2). At Dox 0.5 compared to control, the analysis revealed 43 IR events, 4 A3SS events, and 1 A5SS event that were significantly differentially spliced (Fig. 2C) (S-Table 2). We did not see any significant exon skipping events in our analysis. Out of the statistically significant events we observed maximum overlap of 26 events between Dox 0.1 (vs control) and Dox 0.5 (vs control) (Fig. 2D). There was an overlap of 13 events among all the three comparisons and only 1 event was common between Dox 0.01 (vs control) and Dox 0.5 (vs control) as well as between Dox 0.01 (vs control) and Dox 0.1 (vs control) (Fig. 2D).

### Gene level analysis of *RAP1* mediated alterations in alternative splicing

Next, we investigated the impact of altered *RAP1* expression levels on splicing events for individual genes. Among the various splicing events analyzed, we observed both increased and decreased in intron retention (Percent Spliced Intron (PSI) values) in a monotonic and non-monotonic manner as *RAP1* levels were reduced, underscoring the dual regulatory role of *RAP1* to both increase and decrease the frequency of particular splicing outcomes (Fig. 3A) (S-Table 3). Specifically, 59 splicing events exhibited a monotonic increase in intron retention with decreasing *RAP1* levels, while 14 events showed a monotonic decrease (Fig. 3A) (S-Table 3). Additionally, 121 splicing events displayed a non-monotonic increase in intron retention, and 64 events exhibited a non-monotonic decrease (Fig. 3A). We also observed two splicing events within non-monotonic category with consistent intron retention values at control and Dox 0.5 µg/mL which are classified as “other” (Fig. 3A) (S-Table 3). Therefore, not only does *RAP1* play a dual role (activation or repression) as a transcription factor (62,71,91,92), but it also demonstrates a dual effect in the regulation of IR.

**Fig 3:**
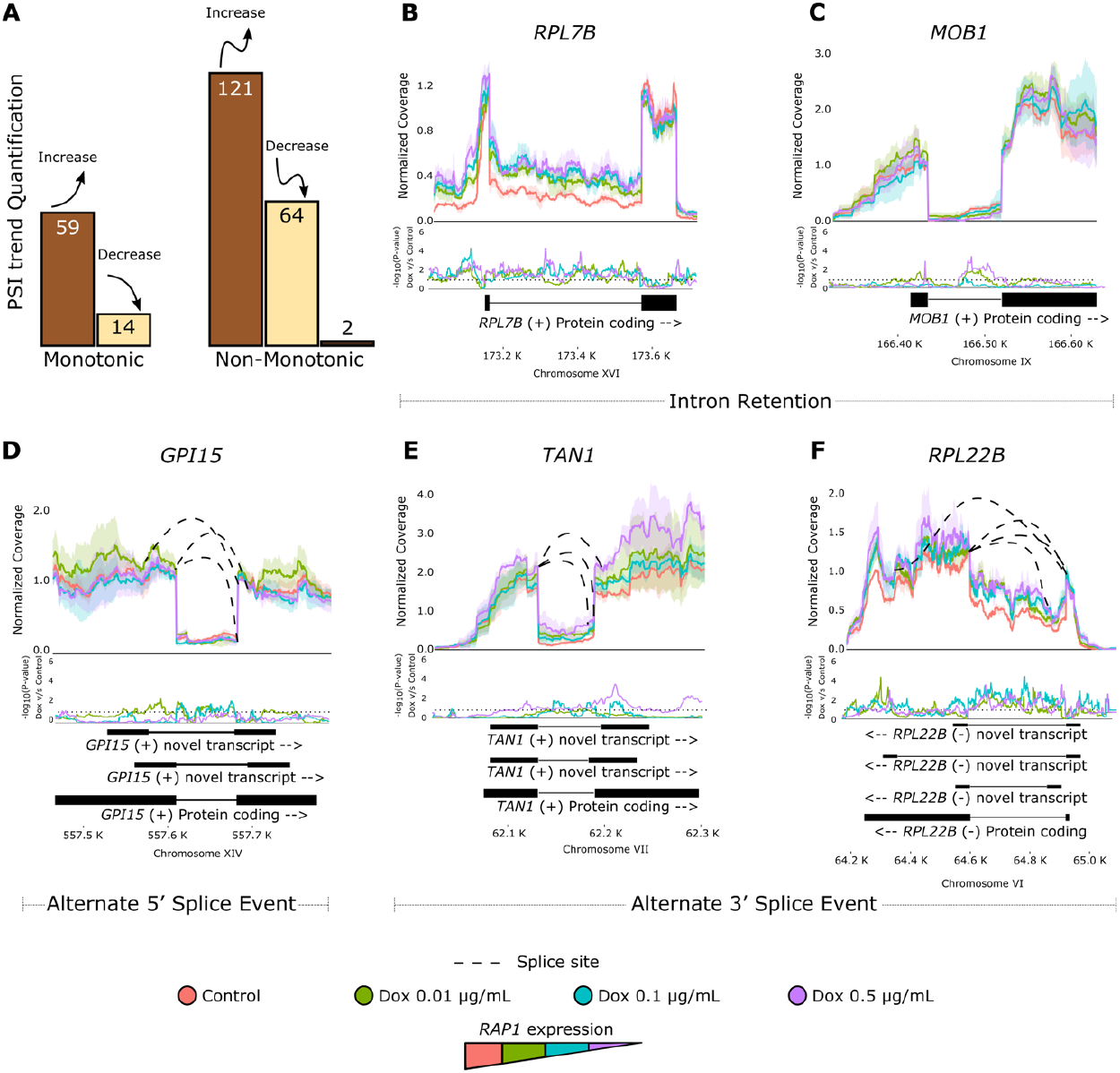
Dual influence of *RAP1* on splicing outcomes. A) Quantification of the effect on Percent Spliced In (PSI) values as *RAP1* levels are reduced. The analysis revealed a dual trend, with both increases and decreases in PSI values observed. These changes occur in monotonic and non-monotonic patterns, emphasizing *RAP1’s* dual role in regulating intron retention. B-F) Coverage plot for B) *RPL7B*, C) *MOB1*, D) *GPI15*, E) *TAN1*, and F) *RPL22B* at varying *RAP1* levels. The top panels display coverage profiles, with dashed lines marking splice sites. The bottom panels show nucleotide-level significance, with dotted lines indicating the significance threshold. Points above the threshold represent significant coverage. Different isoforms are illustrated below the plots, providing a comprehensive view of splicing variation under altered *RAP1* levels.

Representative examples of *RAP1* acting to promote and repress AS are shown in Fig 3. *RAP1* acts as an activator of transcription for Ribosomal 60S subunit 7B (62) (*RPL7B*) (S-Fig. 1A) and we found that reduction of *RAP1* caused significant increases in IR (Fig. 3B). This means *RAP1* acts as an activator of *RPL7B* transcription level and a repressor of *RPL7B* IR. In contrast, we found that *RAP1* acts as a repressor of IR for the Mps one binder (*MOB1*) gene (Fig. 3C) but does not play a role in regulation of transcription level for *MOB1* (S-Fig 1A). In Fig. 3 we also show examples of different splicing events occurring simultaneously on the same gene. For example, for a gene *GPI15* (GlycosylPhosphatidylInositol anchor biosynthesis) we observe A5SS event occurring along with IR. Here we also see reduction in IR as *RAP1* levels are reduced. Even though *RAP1* does not act as a regulator of transcription level (S-Fig. 1A) for *GPI15* we found significant effects of *RAP1* reduction on *GPI15* IR and *GPI15* A5SS splicing (Fig. 3D). We also show in Fig. 3 that the *TAN1* (Trna AcetylatioN) gene has significantly decreased expression level when *RAP1* levels are reduced (therefore *RAP1* acts as a transcriptional activator in wild-type yeast, S-Fig. 1A). At the same Dox concentration, there was a notable increase in IR for *TAN1* (Fig. 3E) as compared to control. *RAP1* reduction also had significant effects on *TAN1* A3SS splicing events, which vary with different *RAP1* dosages (significant at Dox 0.1) (Fig. 3E). Another ribosomal protein gene, *RPL22B* (Ribosomal Protein of the Large subunit) showed a significant decrease in gene expression at all three Dox levels (S-Fig. 1) but also had significantly increased IR at various *RAP1* levels (Fig. 3F). An A3SS splicing event was also observed at different *RAP1* levels, with statistically significant differential splicing at Dox 0.5 observed (Fig. 3F). Overall, we found that *RAP1* does not need to act as a regulator of gene expression level for a particular gene to have an impact on AS for that same gene and some genes have effects from *RAP1* reduction on multiple different types of AS for the same gene.

### Intron retention increases as *RAP1* expression decreases

The majority of splicing events observed in our study were IR (Fig. 2A, C), therefore we wanted to see if there was any pattern associating *RAP1* levels and the amount of IR. Our results indicated that experimentally reduced *RAP1* levels corresponded to a modest but statistically significant increase in IR (Fig. 4A) (S-Table 4). We identified 68 genes exhibiting significant changes in IR at one or more Dox level; 45 of these genes were identified as direct downstream targets of *RAP1* transcriptional regulation (Fig. 4B) (S-Table 5). Therefore, *RAP1* likely plays a role in both directly (as a splicing regulator on specific genes) and indirectly (as a transcription factor regulating expression of splicing factors that then regulate AS) influencing splicing outcomes for genes.

**Fig 4:**
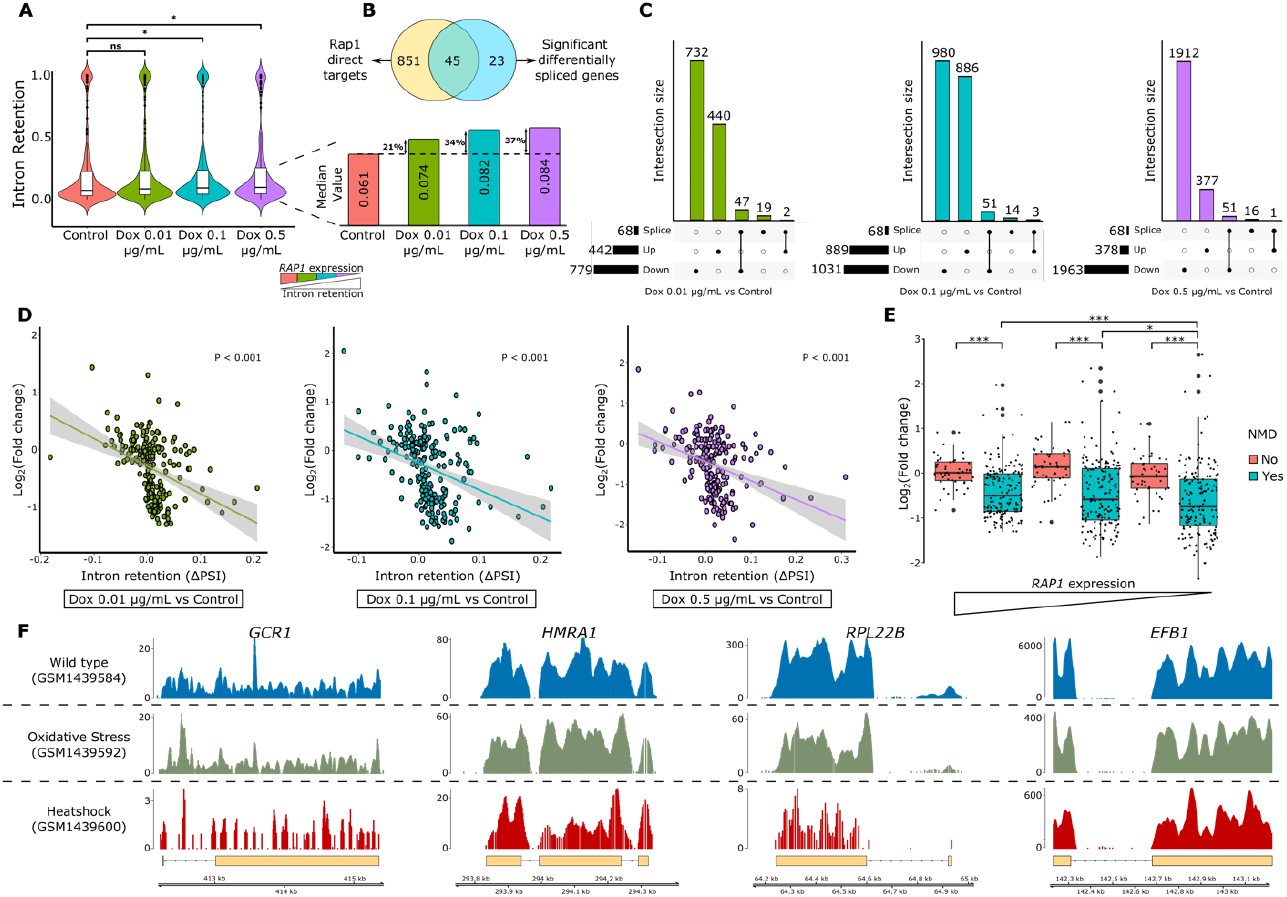
Splicing outcomes and transcriptional changes. A) Quantification of intron retention (y-axis) across various *RAP1* levels (x-axis). A significant increase in intron retention is observed as *RAP1* levels are reduced. Only the splicing events classified under IR were used for this analysis. B) The figure highlights the intersection of Rap1-regulated genes and those showing altered splicing patterns. C) UpSet plots depicting the overlap between significant differentially spliced genes and genes that are differentially regulated (up- or downregulated) at Dox concentrations of 0.01, 0.1, and 0.5 µg/mL. D) Scatter plots visualizing the relationship between Intron retention on x-axis and changes in gene expression (log_2_ fold change) on y-axis at various *RAP1* levels. E) Boxplot showing the change in gene expression (log_2_ fold change) between the genes predicted to undergo nonsense mediated decay (NMD) and those not predicted for NMD across different expression levels of *RAP1*. F) Depiction of ribosomal peaks at intronic and exonic regions under wildtype, oxidative stress, and heat shock conditions. The peak observed at the intronic location indicates that introns are been translated by the ribosomal machinery. *GCR1, HMRA1*, and *RPL22B* all had Riboseq peaks in the intronic sequences, *EFB1*, an intron containing gene that did not show IR in our analysis also didn’t show any ribosomal peaks on its introns in this publicly available riboseq data.

To determine if the genes with significantly differential splicing also exhibited significant changes in gene expression we compared our splicing results to genome-wide gene expression quantification after *RAP1* reduction that was published in our recent study (62). At a Dox concentration of 0.01 (compared to control), we observed an overlap of 47 genes between the 68 significantly differentially spliced genes and the 799 downregulated genes (Fig. 4C), whereas only two genes overlapped between the 68 differentially spliced genes and the 442 upregulated genes (Fig. 4C). At a Dox concentration of 0.1 (compared to control), there was an overlap of 51 genes among the 68 significantly differentially spliced genes and 1,031 downregulated genes (Fig. 4C). At this *RAP1* level, only three genes were both upregulated and significantly differentially spliced (Fig. 4C). When *RAP1* expression was further reduced by using a Dox concentration of 0.5 (compared to control), we found that 51 genes were both significantly differentially spliced and downregulated, while only one gene was both upregulated and significantly differentially spliced (Fig. 4C). These results suggest that reduction in gene expression is associated with differential splicing. We further observed an inverse relationship between *RAP1* expression and intron retention, where higher intron retention correlated with lower *RAP1* levels (Fig. 4A). Furthermore, genes exhibiting significant differential splicing were predominantly downregulated, suggesting a connection between intron retention and gene expression level. Direct comparison revealed that increased intron retention was significantly associated with a reduction in gene expression (Fig. 4D) consistent with a prior study (41).

### mRNAs with retained introns leave the nucleus and are translated by the ribosome

*RAP1* reduction causes altered nucleosome displacement resulting in ectopic transcription initiation within promoters (58) and generation of non-functional or unstable transcripts that are subsequently degraded either in the nucleus by the RNA exosome or in the cytoplasm via the nonsense-mediated decay (NMD) pathway (93–95). Our analysis also identified 67 % of splicing events resulted in the introduction of a PTC and predicted to be degraded by the NMD pathway, even under control conditions (S-Fig 1B). This result is consistent with a previous study focusing on NMD in mutant yeast strains (41) where they also found a large proportion of splicing events in yeast result in a transcript targeted for NMD. Taking a closer look at NMD predicted transcripts and comparing their expression with non-NMD predicted transcripts, we found a significant decrease in gene expression for genes likely targeted for NMD at various *RAP1* levels suggesting that the transcripts of these genes were exported out from the nucleus into the cytoplasm where they were degraded (Fig. 4E). While the number of genes making isoforms with IR remained relatively constant across *RAP1* levels (Fig. S1), the proportion of isoforms per gene that were produced containing retained introns increased (Fig. 4A) and therefore the overall expression level of these genes would be expected to decrease by NMD. Our data were consistent with this hypothesis, genes with transcripts predicted to result in NMD had significantly reduced gene expression than those without predicted NMD (Fig. 4E). This mechanism of gene regulation, where regulated retention of intron resulted in transcripts that contain PTC that are targeted for NMD, which ultimately caused reduced overall gene expression levels has been observed in a few studies (56–59); here we show that *RAP1* plays an important role in gene expression that is regulated with this less common mechanism.

To determine if isoforms with retained introns are indeed exiting the nucleus, we analyzed publicly available RiboSeq data where wild-type yeast was grown under control, oxidative stress, and heat shock conditions (96). We observed ribosome peaks in both intronic and exonic regions of genes, confirming that transcripts with retained introns (e.g. *GCR1, HMRA1, RPL22B)* were exported from the nucleus to the cytoplasm where ribosomes are present indicating that the cytoplasmic NMD pathway is a reasonable mechanism for removal of transcripts with PTC resulting from IR (Fig. 4F). Our results collectively demonstrated that reduced *RAP1* expression increased intron retention (Fig. 4A), sometimes leading to the incorporation of PTC in the transcripts. These transcripts are then exported to the cytoplasm, where they are degraded via the NMD mechanism, ultimately resulting in decreased gene expression (Fig. 4D). This uncovers a new aspect of *RAP1’s* role in gene regulation mediated by splicing.

### Enriched DNA sequence motifs associated with genes undergoing AS in response to *RAP1* reduction

We next asked if there was enrichment of DNA sequence motifs in the intronic sequences of genes with different types of AS. We found enriched motifs for each type of AS. Motifs identified matched known binding motifs for *CAD1, CIN5* and *YAP1* for IR, A3SS and A5SS splicing events (Fig. 5A). When we interrogated differential expression of genes after *RAP1* titration in our prior work (62) we found that both *CAD1* and *YAP1* had significant changes in gene expression at different *RAP1* levels (Fig. 5B). While *CAD1* was significantly upregulated at all Dox levels, *YAP1* was significantly upregulated at higher Dox (Dox 0.01 and Dox 0.1 µg/mL) and therefore lower *RAP1* expression levels (Fig. 5B). The motif enriched for SE events matched known binding motifs for *HCM1* and *STE12*, both of which were significantly downregulated when *RAP1* levels were reduced (Fig. 5B).

**Fig 5:**
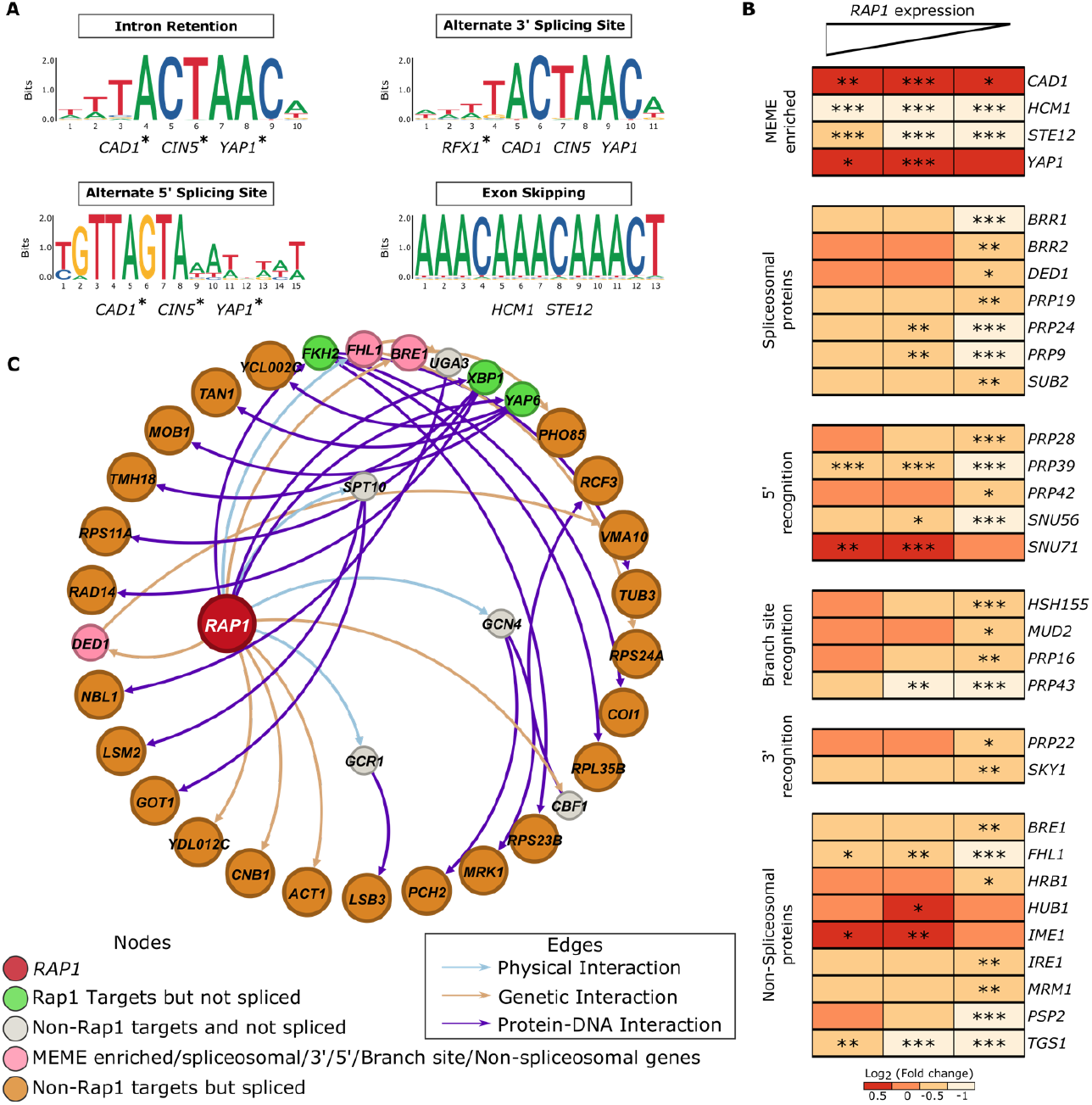
Information flow through *RAP1* interaction network. A) MEME analysis highlighting genes sharing similar enriched motifs for various splicing events. Genes marked with asterisk (*) were identified when only the significant splicing event sequences were used in the analysis while genes without asterisk were identified when all the sequences (regardless of significance) for a specific splicing event were included in the enrichment analysis. B) Heatmap comparing the changes in gene expression (log_2_ foldchange) for genes associated either directly or indirectly with splicing. C) A subsection of genomic network illustrating the information flow from *RAP1* to differentially spliced genes which were not under the direct influence of Rap1 (not direct targets) through intermediated steps highlighting key interactions between different nodes.

### information flow through a genetic network

For efficient splicing to occur, the spliceosomal protein complex must assemble on pre-mRNA to catalyze the precise removal of introns and joining of exonic sequences. This process depends on proteins that recognize intron boundaries at the 5′ and 3′ splice sites, as well as the branch point site. Any lack of nucleotide-level precision during intron removal can impact splicing efficiency, potentially altering the resulting coding sequence and therefore protein function (12). Beyond spliceosomal components, other non-spliceosomal proteins interact with pre-mRNA or spliceosomal elements to modulate splicing activity (97). These proteins can act as splicing enhancers or inhibitors, promoting or blocking splicing at specific sites. Consequently, changes in these regulatory proteins can shift splicing outcomes, influencing which transcript isoforms are produced and ultimately affecting gene expression and cellular function. Our RNA-seq analysis (62) revealed that reducing *RAP1* levels led to a statistically significant decrease in the expression of numerous genes encoding spliceosomal protein complex proteins (Fig. 5B). A similar reduction in gene expression was observed for genes responsible for recognizing the 5′ and 3′ splice sites and the branch point (except for *SNU71*) (Fig. 5B). Additionally, genes encoding non-spliceosomal proteins involved in splicing regulation also had significantly decreased expression as *RAP1* levels were reduced with only *HUB1* and *IME1* significantly increased at certain Dox levels (Fig. 5B). These findings suggest that *RAP1* plays a critical role in regulating the expression of splicing-related genes, thereby indirectly influencing splicing outcomes in yeast.

To understand how *RAP1* signaling spreads in the genome and influences splicing outcomes, a subsection of the genomic network was created (S-Fig. 2)(S-Table 6). This network consists of all *RAP1* protein-DNA targets as well as *RAP1’s* other interactions, including protein-protein and protein-RNA associations. It also includes targets of genes enriched in splicing events (IR, A3SS, A5SS, and SE) identified through MEME analysis (98), regulators of the 23 genes that were significantly differentially spliced at varying *RAP1* levels that are not direct *RAP1* targets (Fig. 4B), and targets of genes associated with spliceosomal components along with genes involved in recognizing the 5′ and 3′ splice sites and branch points (Fig. 5B). Additionally, the targets of non-spliceosomal genes (Fig. 5B) relevant to splicing regulation were also incorporated. The resulting directed network comprises 5,254 nodes and 31,177 edges (S-Fig. 2), providing a comprehensive view of potential regulatory interactions influencing splicing in the context of *RAP1* regulation.

Our analysis reveals both direct and indirect influences of *RAP1* levels on splicing outcomes and gene regulation. Some of the examples are discussed below. Directly, *RAP1* affects the splicing of 45 genes (Fig. 4B). Indirectly, *RAP1* affects intermediary genes such as *FKH2* through protein-DNA interactions, which subsequently impacts downstream targets like *TUB3* (Fig. 5C). *RAP1* also exerts regulatory influence through its physical association with *FHL1*, a non-spliceosomal gene. By altering *FHL1* expression, *RAP1* can indirectly affect splicing through a multi-step pathway (*RAP1* → *FHL1* → *UGA3* → *NBL1*) (Fig. 5B). This comprehensive network highlights the complexity of *RAP1’s* role in splicing regulation and gene expression (S-Table 7).

## Discussion

Our analysis of the of the role of *RAP1* in regulating mRNA splicing provides new insights into the multifaceted roles of this well-studied transcription factor. In our previous study, we showed that by utilizing the Tet-off system one can titrate the expression of *RAP1* by adding different concentrations of Dox in growth media (62). Using the same approach, in this study we assessed the impact of various *RAP1* levels on alternative splicing in yeast using RNA sequencing (Fig. 1A, B). Our analysis revealed the majority of alternative splicing events are intron retention while only a small proportion belonged to A3SS and A5SS (Fig. 2A, C). We also found that as *RAP1* levels decreased, a notable increase in IR was observed (Fig. 4A).

Gene level analysis identified specific genes such as *RPL7B*, a downstream target of *RAP1*, was directly influenced by changes in *RAP1* expression in terms of splicing and gene expression (Fig. 3C). We saw a significant increase in IR for this gene as *RAP1* expression was reduced. On the other hand, *RAP1* showed an opposite effect on IR in *MOB1* as well as *GPI15* gene where intron retention was reduced as *RAP1* levels were reduced (Fig. 3D, E). Overall, we observed 180 splicing events where PSI values increased and 78 splicing events where PSI values decreased as *RAP1* levels were reduced. Thus, our results demonstrate *RAP1’s* dual regulatory role as an activator and repressor in both transcription and splicing.

This relationship between *RAP1* levels and splicing outcomes was further shown when 45 out of the 68 genes that were significantly differentially spliced were also direct transcriptional regulatory targets of *RAP1* (Fig. 4B). Comparing our results with our previous gene expression data, we observed a substantial overlap between significantly downregulated genes and significantly differentially spliced genes (Fig. 4C). To delve deeper into indirect effects through the regulatory network influenced by *RAP1*, we constructed a genomic network (S-Fig. 2) which allowed us to map a comprehensive network reflecting both the direct and indirect regulatory influence of *RAP1*. A decrease in *RAP1* expression results in a significant reduction in the expression of genes associated with the spliceosome, as well as those involved in recognizing critical intron sites, including the 5′ and 3′ splice sites and the branch point (Fig. 5B). Additionally, some non-spliceosomal genes show a similar decrease in expression (Fig. 5B). This reduction in expression of these critical genes can impact splicing efficiency, potentially leading to the production of altered transcript isoforms. Changes in *RAP1* levels propagate through the network in a stepwise manner via protein-DNA interactions, as well as physical and genetic interactions between genes, highlighting *RAP1’s* broader regulatory influence on splicing mechanisms, direct and indirectly. Our analysis identified candidate regulatory proteins/genes forming a potential pathway by which *RAP1* may indirectly affect splicing processes (Fig. 5C).

In this study, we also noted an increase in IR correlated with reduced *RAP1* levels. Furthermore, this increase in intron retention was accompanied by a corresponding decrease in gene expression, indicating an inverse relationship between intron retention and gene expression levels as *RAP1* expression diminished (Fig. 4D). This suggests that *RAP1* not only regulates gene expression directly but also modulates splicing events, particularly intron retention, which in turn impacts overall gene expression. We also observed that intron-retained isoforms are exported out of the nucleus with ribo-seq data, but our data suggest that many of these transcripts undergo mRNA degradation via the NMD pathway (Fig. 4E, F) because a PTC was introduced during intron retention. As the majority intron-containing genes are ribosomal protein genes and *RAP1* acts as an activator of these RP genes, our analysis explains one possible mechanism why a decrease in *RAP1* causes a reduction in the expression of ribosomal protein genes.

Overall, our study provides important new insights into the regulatory role of *RAP1* in mRNA splicing. It not only uncovers previously uncharacterized functions of *RAP1* in splicing but also suggests that *RAP1* may play an important role in regulating genes through the less common IR-PTC-NMD mechanism resulting in downregulation of gene expression. Additionally, *RAP1’s* conserved functions between yeast and humans make yeast an excellent model to explore its molecular mechanisms. Investigating *RAP1’s* role in splicing in yeast could help uncover conserved pathways and clarify whether *TERF2IP* directly or indirectly influences splicing in humans potentially leading to new understanding of diseases associated with *TERF2IP*.

## Materials and Methods

### Sample Preparation

In this study, the Tet0_7_-*RAP1 Saccharomyces cerevisiae* strain derived from the R1158 control background strain was utilized. This strain features a tetracycline response element upstream of *RAP1* and expresses the Tetracycline-controlled Trans Activator (tTa) protein (99). Following inoculation of a colony of this strain into overnight cultures, subsequent dilutions were made the following day, and cultures were grown until reaching the exponential phase (OD 0.6-0.8). At this point, cultures were once again diluted, and *RAP1* levels were modulated by the addition of Dox to the culture media (Fig. 1A). In the absence of Dox, the Trans Activator protein interacts with the tetracycline response element, resulting in maximum *RAP1* expression. Conversely, the presence of Dox induces alterations in *RAP1* levels, with higher Dox concentrations corresponding to lower *RAP1* expression levels. Further elaboration on this methodology can be found in our previous research (62). Upon reaching the exponential phase, cultures were centrifuged, and pellets were collected and processed for RNA extraction.

### RNA extraction

Yeast pellets, exhibiting various *RAP1* levels (Dox concentrations: 0 µg/mL (control), 0.01 µg/mL, 0.1 µg/mL, and 0.5 µg/mL), underwent initial treatment with 20T lyticase for 30 minutes at 30°C. Subsequently, RNA extractions were carried out utilizing the Promega SV Total RNA isolation kit using a modified protocol (100). Assessment of RNA quality was performed using a Nanodrop spectrophotometer, followed by the submission of samples to the Michigan Sequencing Core for sequencing on an Illumina Hiseq 4000 platform. Notably, three replicates were sequenced for each sample to ensure robustness and reliability of the data.

### Read Alignment and splicing quantification

The Fastq files obtained from the sequencing core underwent quality assessment using the FastQC tool (87) within the Galaxy platform (101). Subsequently, the reads were aligned to the *Saccharomyces cerevisiae* reference genome obtained from the Ensembl genome browser using the STAR aligner (88) tool with default parameters. The resulting BAM files were then processed through SpliceWiz (89), an R-based package utilized for the quantification of splicing events, differential splicing analysis across various conditions, and the generation of coverage maps for visualization purposes.

### Riboseq Analysis

To investigate whether intron-retained transcripts reach the translational machinery, we analyzed publicly available Ribo-Seq data generated under wild type (GSM1439584), oxidative stress (GSM1439592), and heat shock (GSM1439600) conditions (96). The quality check of these fastq files obtained from NCBI was performed in Galaxy server using FastQC tool (87). Subsequent alignment to the *Saccharomyces cerevisiae* reference genome (obtained from Ensembl) was conducted using the STAR aligner (88) on a high-performance computing (HPC) cluster. The resulting BAM files were further analyzed and visualized utilizing the Gviz (102) and GenomicRanges (103) libraries in R. Representative examples are shown in Fig 4F.

### Motif enrichment and network construction

The coordinates of the splicing events under various *RAP1* levels were used to extract nucleotide sequences through BioMart (104). These sequences, corresponding to each splicing event category (IR, A3SS, A5SS, and SE), were uploaded to the MEME Suite (98). Enrichment analysis was performed using XSTREME (105) with the YEASTRACT (106) database as the background. The analysis was conducted twice: first, using only sequences from significant splicing events (indicated by asterisks on those genes), and second, using all sequences regardless of their significance level.

For network construction, we compiled a comprehensive list of relevant gene interactions to capture the full regulatory landscape of *RAP1*-influenced splicing events. This list included: (a) *RAP1* targets, encompassing both direct protein-DNA and protein-protein interactions, as well as protein-RNA interactions retrieved SGD (107) (b) protein-DNA targets of 45 genes that were significantly differentially spliced and were downstream targets of *RAP1* (Fig. 5B) (c) targets of genes identified through MEME enrichment analysis, along with genes involved in critical spliceosome functions—specifically, those responsible for recognizing and binding the 5′ splice site, 3′ splice site, and branch sites of introns, as well as associated spliceosomal components that facilitate spliceosome assembly and stability along with some non-spliceosomal associated proteins (Fig. 5B) (d) regulators of the 23 genes (extracted from SGD (107)), that were significantly spliced at various *RAP1* levels but were not the direct targets of *RAP1* (Fig. 4B). This list was used as input to generate the network using Gephi (108). The shortest path from *RAP1* to these 23 spliced genes were computed using “Dijkstra algorithm” in R package “igraph” (109).

## Supplementary Figure Legends

S-Fig 1: A) The values in the heatmap represent the log_2_ foldchange with significant changes highlighted by asterisk (*) at various *RAP1* levels. B) The plot comparing the frequency of splicing events predicted to undergo NMD with those not predicted for NMD across different conditions.

S-Fig 2: Interaction network highlighting information flow from *RAP1* to different genes (nodes) highlighting key interactions between different nodes.

## Supplementary Tables

S-Table 1: Splicing events observed at various *RAP1* (Dox) levels.

S-Table 2: List of differential splicing events observed when Dox levels (0.01,0.1, and 0.5 µg/mL) were compared to control (Dox 0).

S-Table 3: AvgPSI trends across various *RAP1* levels.

S-Table 4: AvgPSI trends exclusive to IR events across various *RAP1* levels.

S-Table 5: List of overlaps between *RAP1* targets and significant differentially spliced genes.

S-Table 6: Interaction file for *RAP1* network.

S-Table 7: Path from *RAP1* to 23 significantly differentially spliced genes.

## Acknowledgements

Research reported in this publication was supported by Wesleyan University (start-up funds to J.D.C. and Department of Biology funds to J.D.C.) and the National Institute of General Medical Sciences of the National Institutes of Health under Award Number R15GM135901 (awarded to J.D.C.). The content is solely the responsibility of the authors and does not necessarily represent the official views of the National Institutes of Health.

## Data Deposition

Gene expression data used in this manuscript are deposited in the Gene Expression Omnibus (GEO) under accession number GSE226065.

